# Effects of Simvastatin on RBL-2H3 cell degranulation

**DOI:** 10.1101/2022.09.25.509437

**Authors:** Michiko Yoshii, Ai Kitazaki, Koichiro Ozawa

**Affiliations:** Department of Pharmacotherapy, Graduate School of Biomedical and Health Sciences, Hiroshima University, 1-2-3, Kasumi, Minami-ku,Hiroshima 734-8553,Japan; School of Pharmaceutical Sciences, Hiroshima University; 1-2-3, Kasumi, Minami-ku,Hiroshima 734-8553,Japan

**Keywords:** Simvastatin, rat basophilic leukemia-2H3 cell, degranulation, geranylgeranyl pyrophosphates, protein kinase C delta

## Abstract

Hypercholesterolemia is a major complication of arteriosclerosis. Mast cells in arteriosclerosis plaques induce inflammatory reactions, and promote arterial sclerosis. In this study, we evaluated the pharmacological effects of simvastatin (SV)-3-hydroxy-3-methylglutaryl-CoA (HMG-CoA) reductase inhibitors on the degranulation of rat basophilic leukemia (RBL)-2H3 cells, which are commonly used as mast cell models. SV significantly decreased degranulation induced by three types of stimulation: antigen antibody reaction (Ag-Ab), thapsigargin (Tg) serosal endoplasmic reticulum calcium ATPase (SERCA) inhibitor, and A23187 calcium ionophore. SV had a stronger inhibitory effect on degranulation induced by Ag-Ab stimulation than the other two stimulations. However, SV had no effect on increases of intracellular Ca^2+^ concentrations. Mevalonate or geranylgeraniol co-treatment with SV completely prevented the inhibitory effect of SV on degranulation induced by these stimulations. Immunoblotting results showed that SV inhibited protein kinase C (PKC) delta translocation that was induced by Ag-Ab but not by Tg or A23187. SV induced a reduction in active Rac1, and actin filament rearrangements.

In conclusion, SV inhibits RBL-2H3 cell degranulation by inhibiting downstream signaling pathways, including the sequential degranulation pathway. These inhibitory effects were completely reversed by the addition of geranylgeraniol and might be induced by changes in the translocation of the small GTPase families Rab and Ras and Rho, which are related to vesicular transport and PKC delta activation, respectively. These changes are caused by the inhibition of HMG-CoA reductase by SV following the synthesis of farnesyl and geranylgeranyl pyrophosphates, which play important roles in the activation of small GTPases.

## Introduction

The statin SV is an inhibitor of HMG-CoA reductase, which is a rate-limiting enzyme of the cholesterol biosynthesis pathway. Statins are effective drugs for hypercholesterolemia, which causes myocardial infarction, cardiac angina, and brain infarction, and the cholesterol-lowering effect of statins has reduced the risk of cardiovascular disease (CVD) in large, randomized studies in recent decades.

Intraplaque mast cell numbers are associated with plaque vulnerability, including the lipid core size, intraplaque hemorrhage, microvessel density, and inflammatory cell accumulation, and they are also correlated with future cardiovascular events. Therefore, mast cells actively contribute to atherosclerotic plaque progression and destabilization ^1,2)^. Exocytosis of mast cells plays an important role in mediating allergic and inflammatory pathogenesis and releases numerous inflammatory compounds, such as histamine, lysosomal compound, and β-hexosaminidase ^3)^.

The effects of statins on CVD has led to additional concerns regarding the pleiotropic effects of statins^4)^. A multicenter fluorodeoxyglucose (FDG)-positron emission tomography/computed tomography feasibility study by Tawakol et al. showed that statins decrease vascular inflammation in patients at an increased risk of CVD. Statins induce significant rapid dose-dependent reductions in FDG uptake, which likely represent changes in atherosclerotic plaque inflammation^5^). The mevalonate pathway, which is inhibited by statins, is important for the production of cholesterol as well as intermediates, such as farnesyl pyrophosphate and geranylgeranyl pyrophosphate, which are used in the modification of the small GTPase families Rab, Arf, Ras, and Rho for translocation^6,7^). Incomplete modification of these signaling molecules by statins likely induces a decrease in signal transduction and causes pleiotropic effects. The Rab family binds geranylgeranyl pyrophosphate via geranylgeranyl pyrophosphate transferase I and II, which is related to vesicular transport and docking to the plasma membrane^8)^.

However, the molecular mechanisms responsible for the inhibitory effects of statins on degranulation have not yet been elucidated. Antigen cross-linking IgE bound to FcεR1 activates mast cells and LYN-dependent phosphorylation of immunoreceptor tyrosine-based activation motifs (ITAMs) and leads to the additional activation of tyrosine-protein kinases (FYN and SYK). These activations lead to the phosphorylation of adaptor proteins, such as linker for activation of T cells (LAT) and GRB2-associated binding protein 2 (GAB2). LAT activates phospholipase Cγ (PLCγ), which generates inositol-1,4,5-trisphosphate (InsP3) and diacylglycerol (DAG), leading to an increase in intracellular Ca^2+^ concentration, followed by the activation of PKC alpha, beta, and gamma. GAB2 activates phosphoinositide 3-kinase (PI3K) that generates phosphatidylinositol-3,4,5-trisphosphate (PtdIns(3,4,5)P3), thus leading to the activation of PKCs, including PKC delta. PI3K activity is induced and reinforced by a distributed network of the small GTPase families Ras and Rho^9)^. The increase in intracellular Ca^2+^ levels and the activation of PKCs trigger the sequential degranulation process, secretory vesicle translocation to the plasma membrane, docking vesicle, and plasma membrane, and the release of contents in vesicles. For these process, small GTPases, especially the Rab family, are important^3,8,10^). Thirty Rab proteins are related to the exocytosis of mast cells^11^), and Sahid et al. suggested that SV decreases the complex of Rab27a and Doc2a to promote exocytosis by suppressing the geranylgeranylation of Rab27a^12)^.

Previous studies indicate that PKC delta plays an essential role in degranulation, although all isoforms of PKC in RBL-2H3 cells contain activated isoforms of PKC, PKC alpha, beta, delta, epsilon, and zeta. PKC beta and delta at sufficient concentrations represent potent transducers of signals for exocytosis in antigen-stimulated permeabilized cells ^13–15)^. PKC delta plays an essential downstream mediating role in the degranulation elicited by Ca^2+^ mobilization, and reactive oxygen species mediate the activation of PKC delta by Ca^2+^ in the regulation of degranulation ^16)^. PKC delta is required to switch tomosyn-1 from an inhibited form to an interacting form that partners with syntaxin (STX) during granule fusion ^17)^.

Actin rearrangement is related to degranulation associated with the Rho small GTPases RhoA, Rac1, and CDC42 ^18)^. Farnesyl and geranylgeranyl pyrophosphates play important roles in the translocation of Rho small GTPases. A recent study reported that tuning cytoskeletal actin dynamics may represent a process by which RBL cells can efficiently coordinate their activation ^19)^. However, whether exocytosis of mast cells is relevant to actin rearrangement remains controversial ^20)^.

In this study, we investigated the effect of SV on the degranulation of RBL-2H3 cells stimulated with Ag-Ab, Tg, or A23187. SV does not inhibit the increase in [Ca^2+^] i associated with any stimulation. However, SV does inhibit mast cell degranulation via depletion of intracellular geranylgeraniol PP, which is essential for the activation of small GTPases, because the inhibitory effect of SV was fully recovered by mevalonic acid and geranylgeraniol but not by farnesol. SV decreases PKC delta activation induced by Ag-Ab stimulation and alters the actin filament rearrangement. These additional effects of SV might be to inhibit the degranulation of RBL-2H3 cells upon stimulation.

## Materials and Methods

### Chemicals

The following chemicals were used: RPMI-1640 powder (High Clone, USA); FBS (Thermo Fisher, USA); thapsigargin (FujifilmWako Chemicals, Japan); A23187 (Cayman Chemicals, USA); mouse monoclonal anti-STIM1 (Abnova, Taiwan), Fluo8/AM (AAT Bioquest, USA); and anti-DNP IgE, TritonX-100, p-nitrophenyl-N-acetyl-β-D-glucosaminide, and DNP (dinitrophenyl)-BSA (Sigma-Aldrich, Japan).

### RBL-2H3 cells

Rat basophilic leukemia (RBL)-2H3 cells were maintained in RPMI-1640 supplemented with 15% FBS, 100 U/ml penicillin, 100 μg/ml streptomycin, 250 ng/mL amphotericin, and 2 mM glutamine. The cells were sensitized overnight with 0.5 mg/ml anti-dinitrophenol IgE (anti-DNP IgE) before use. The cells were simultaneously pretreated for 24 h with SV at 0–150 nM.

### Degranulation

We measured the release of β-hexosaminidase as an index of mast cell degranulation using a previously described method ^21)^. Briefly, cells were seeded at 4×10^4^ cells/well, sensitized with anti-DNP-IgE, and treated with SV for 24 h. The cells were washed with Siraganian buffer (125.4 mM NaCl, 11.5 mM glucose, 5.9 mM KCl, 1.2 mM MgCl_2_, 1 mM CaCl_2_, 10 mM HEPES, 0.1% BSA; pH 7.4), and then IgE-sensitized RBL-2H3 cells were stimulated with 20 ng/mL dinitrophenol-bovine serum albumin (DNP-BSA) at 37 °C for 15 min. Non-sensitized cells were stimulated for 15 min with Tg (2.0 μM) or A23187 (0.5 μM). To test the restorative ability of SV, cells were treated with 300 μM mevalonolactone, 20 μM geranylgeraniol, and 20 μM farnesol. Triton X-100 solution (0.2%) was added to the cells as a positive control. The supernatant (10 μL) was then transferred to a 96-well plate. To each well, 10 μL of 1.3 mg/ml p-nitrophenyl-N-acetyl-β-D-glucosaminide in 0.04 M sodium citrate (pH 4.5) was added. The plates were incubated at 37 °C for 1 h, and then 250 μL 0.2 M carbonate-bicarbonate buffer (pH 10.0) was added to each well. The absorbance at 415 nm with a reference filter of 600 nm was measured using a plate reader (Thermo Fisher Scientific, Tokyo, Japan). The percentage of degranulation was calculated using the following formula: % degranulation = OD_415-600nm_/OD_Triton X-100 415-600nm_×100.

### Measurement of the intracellular calcium concentration ([Ca^2+^]_i_)

[Ca^2+^]_i_ was measured using Fluo8/AM. RBL-2H3 cells pretreated with SV for 24 h on glass bottom dishes were loaded with 1 μM Fluo8/AM in Siraganian buffer with 1 mM CaCl_2_ or 0.2 mM EGTA for 30 min at 37 °C and 95% CO_2_. Every 10 s, the fluorescence of an image was captured at 488 nm using a fluorescence microscope (BIOREBO 9000, Keyence, Japan).

### Western blotting for aggregated or monomeric STIM1

After stimulation, the RBL-2H3 cells samples were prepared for Blue Native(BN)-Page and SDS-Page by the procedure of Wittig I. et al. ^22)^. Cells were extracted with 0.8% DDM in solubilization buffer (50 mM Sodium chloride, 50 mM imidazole/HCl, 2 mM 6-Aminohexanoic acid, 1 mM EDTA, pH7.0) following harvest. Harvested samples were centrifuged at 15,000rpm for 30 min. Supernatant (35 μL) + 50% glycerol (5 μL) + 5% CBB (1 μL) were loaded and resolved via BN-PAGE using 4-16% Bis-Tris gels for the analysis of aggregated STIM1. Then, 1/3 volume sample buffer was added to the remaining supernatant, which was boiled for 3 min, loaded, and resolved by SDS-PAGE using 10% acrylamide gel for the analysis of monomeric STIM1. After electrophoresis, the separated proteins were transferred onto polyvinylidene difluoride membranes. After blocking by 5% skim milk, anti-STIM1 antibody was used at a dilution of 1:200 and secondary mouse antibody was used at a dilution of 1:20,000.

### Western blotting for membrane-associated protein PKC

Membrane associated proteins were collected by the procedure of Taguchi Y. et al. ^23)^. Briefly, RBL-2H3cells were detached after incubation for a few minutes in 3 mM EDTA PBS. The collected detached cells were centrifuged at 1,000×g for 5 min, and then the cell pellet was added to phosphate-buffered 2% TX114 lysis buffer, vortexed for 30 s, and then incubated on ice for 30 min. After cell debris was removed by centrifugation, TX114 lysate was incubated at 37 °C for 10 min following centrifugation at 22,500×g for 10 min at RT for phase separation. After removal of the aqueous phase, an equal volume of methanol:chloroform (1:4) was added to the remaining detergent phase, which was incubated on ice for 20 min and centrifuged at 16,000×g for 30 min at 4 °C. After removal of the upper phase, the remaining lower phase was added to methanol in a 9-fold volume and then centrifuged at 16,000×g for 30 min at 4 °C. After removing the supernatant, the remaining pellet was added to 1× sample buffer and boiled prior to analysis. The samples were loaded and resolved via SDS-PAGE using 10% acrylamide gel and then transferred to polyvinylidene difluoride. After blocking with 5% skim milk, anti-PKC delta antibody was used at a dilution of 1:200 and secondary rabbit antibody was used at a dilution of 1:20,000.

### Fluorescence staining of F-actin

To visualize the filamentous actin, RBL-2H3 cells grown on glass coverslips were rinsed with PBS at room temperature and then were fixed with 4 % paraformaldehyde in PBS for 10 min at RT. The fixed cells were permeabilized with 70% ethanol for 5 min and then incubated with phalloidin rhodamine (Molecular Probes, Eugene, OR, U.S.A.) for 60 min at room temperature. Images were captured using a fluorescence imaging microscope (BIOREVO 9000, Keyence, Tokyo, Japan) equipped with excitation (490 nm) and emission (520 nm) filters.

### Pull-down assay for active Rac1

RBL-2H3 cells were lysed on a culture plate with 1 mL of lysis/binding/wash buffer. Clarified cell lysate (500 μg) was treated with either GTPγS (positive control) or GDP (negative control). Half of the non-treated lysates from the SV-treated RBL-2H3 cells and control RBL-2H3 cells and the treated lysates were incubated with 20 μg GST-Pak1-PBD (for active Rac1) (Cell Biolabs, Inc., USA). Each elution and the remaining half of the lysates were analyzed by SDS-PAGE and detected by western blotting using Rac1 primary antibody.

### Statistical analysis

The results of the experiments are expressed as the mean ± S.E. ANOVA followed by Dunnett’s test was used for comparisons between two or more groups. Statistical significance was set at p<0.05.

## Results

### SV decreases mast cell degranulation by Ag-Ab, Tg, and A23187 stimulation

We examined the effect of SV on stimulator-induced degranulation of RBL-2H3 cells. For Ag-Ab stimulation, RBL-2H3 cells sensitized by anti-DNP-IgE (0.5 μg/ml) and pretreated with SV at 50, 75, 100, and 150 nM for 24 h were stimulated with 20 ng/ml DNP-BSA. For Tg or A23187 stimulation, non-sensitized RBL-2H3 cells pretreated with SV in the same manner were stimulated with 2 μM Tg or 500 nM A23187. Values represent the percentage of maximum degranulation (Fig. 1). SV significantly inhibits degranulation of RBL-2H3 cells stimulated by the three kinds of stimulation in a concentration-dependent manner. SV inhibited the degradation induced by DNP-BSA with slightly greater potency than that induced by the other two stimulations. Moreover, the pretreatment did not affect the amount of β-hexosaminidase in the cells (data not shown).

**Fig. 1.**
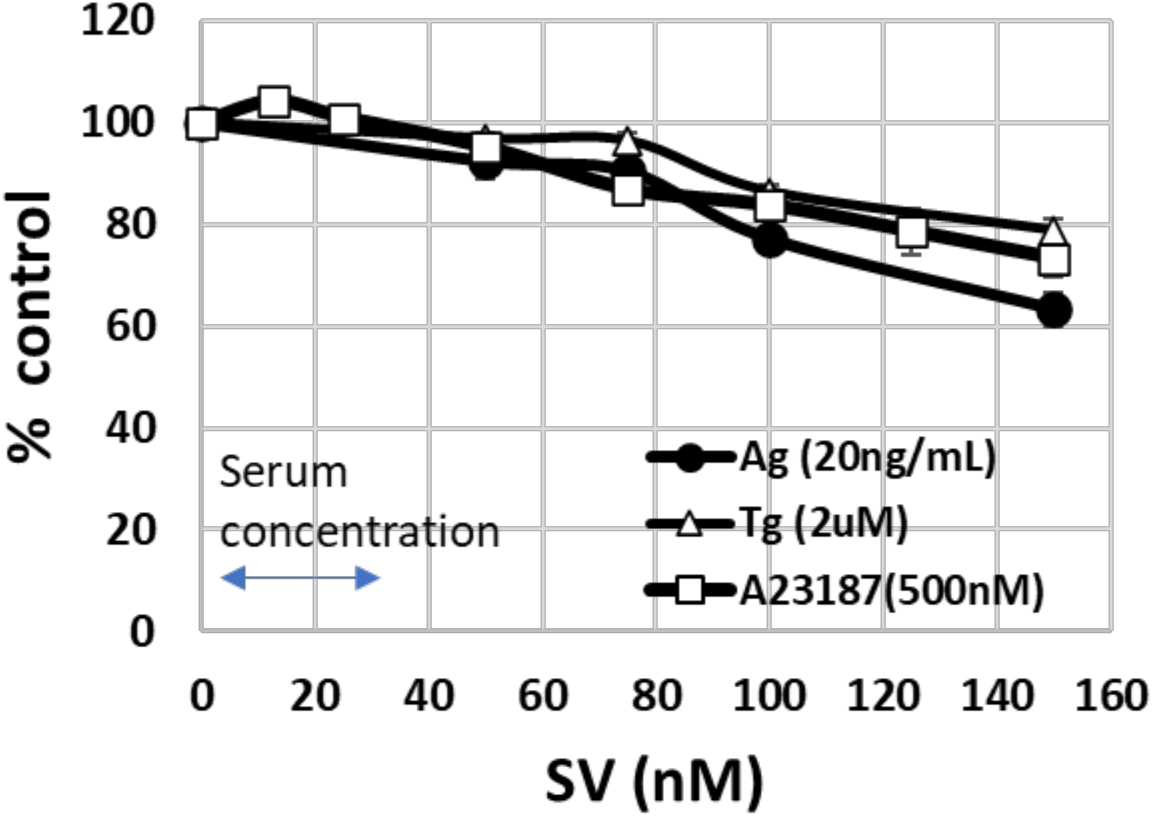
SV pretreatment of RBL-2H3 cells does not inhibit the degranulation by Ag-Ab, Tg, or A23187 stimulation. RBL-2H3 cells pre-treated with SV for 24 h were stimulated for 15 min with DNP-BSA (20 ng/mL), thapsigargin (2 μM), or A23187 (500 nM) in Siraganian buffer. The closed circle, open triangle, and open square lines represent Ag-Ab, Tg, and A23187, respectively. All degranulation values are represented on an index of 100, with 100 representing the % of degranulation from 0 nM SV-pretreated cells. *: p<0.05, **: p<0.01 vs SV 0 nM

### SV does not influence increased [Ca^2+^]_i_ by Ag-Ab, Tg or A23187 stimulation

The increase in [Ca^2+^]_i_ is essential for mast cell degranulation; therefore, we investigated whether applying the SV pretreatment for 24 h increased the [Ca^2+^]_i_ in cells stimulated with Ag-Ab, Tg, or A23187. The [Ca^2+^]_i_ increase was not affected by the SV pretreatment (Fig. 2 a-c).

**Fig. 2.**
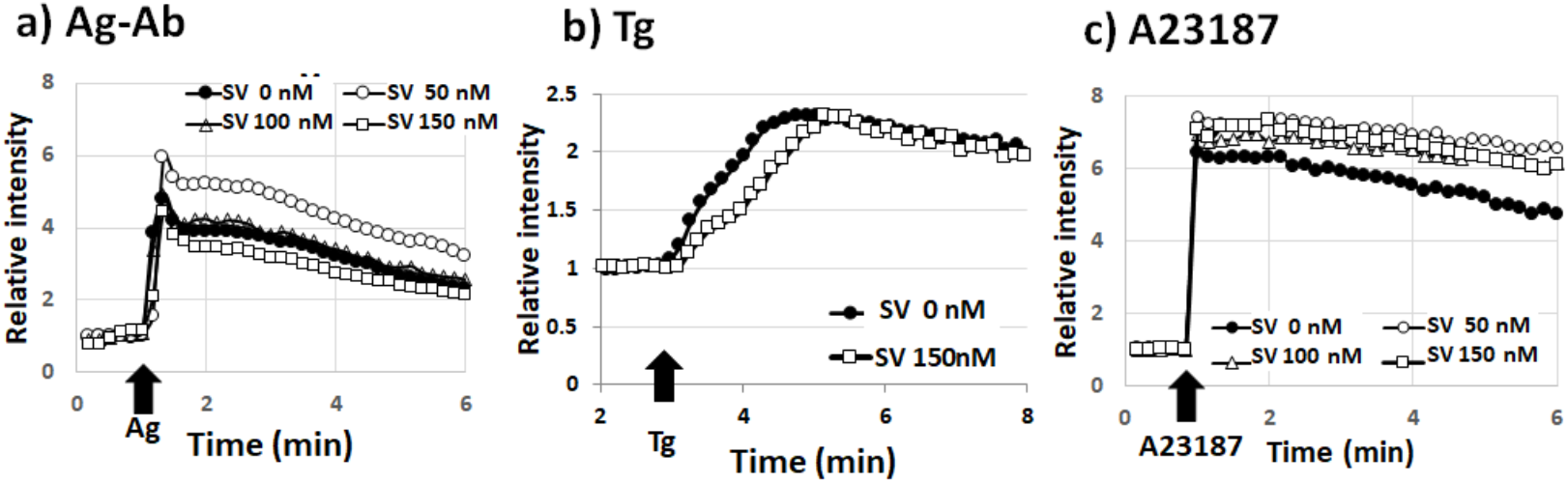
SV pretreatment does not inhibit the increase of [Ca^2+^]_i_ by Ag-Ab or Tg or A23187 stimulation. RBL-2H3 cells pretreated with SV for 24 h were loaded with 1 μM Fluo8/AM for 30 min and then stimulated with) DNP-BSA (20 ng/mL), b) thapsigargin (1 μM), or c) A23187 (500 nM) in Siraganian buffer. The fluorescence change over time was detected under a fluorescence microscope (BIOREVO 9000, Keyence, Japan). The closed circle, open circle, open triangle, and open square lines represent SV at 0, 50, 100 nM, and 150 nM, respectively.

The increase in [Ca^2+^]_i_ associated with Ag-Ab stimulation consists of Ca^2+^ efflux from the endoplasmic reticulum (ER), followed by extracellular Ca^2+^ influx. Therefore, we investigated whether applying the SV pretreatment for 24 h increased the [Ca^2+^]_i_ of cells stimulated with Ag-Ab under extracellular Ca^2+^ deficient conditions caused by EGTA 0.2 mM. The SV pretreatment did not affect the transient increase in [Ca^2+^]_i_ caused by Ca^2+^ efflux from the ER (Fig. 3a).

**Fig. 3.**
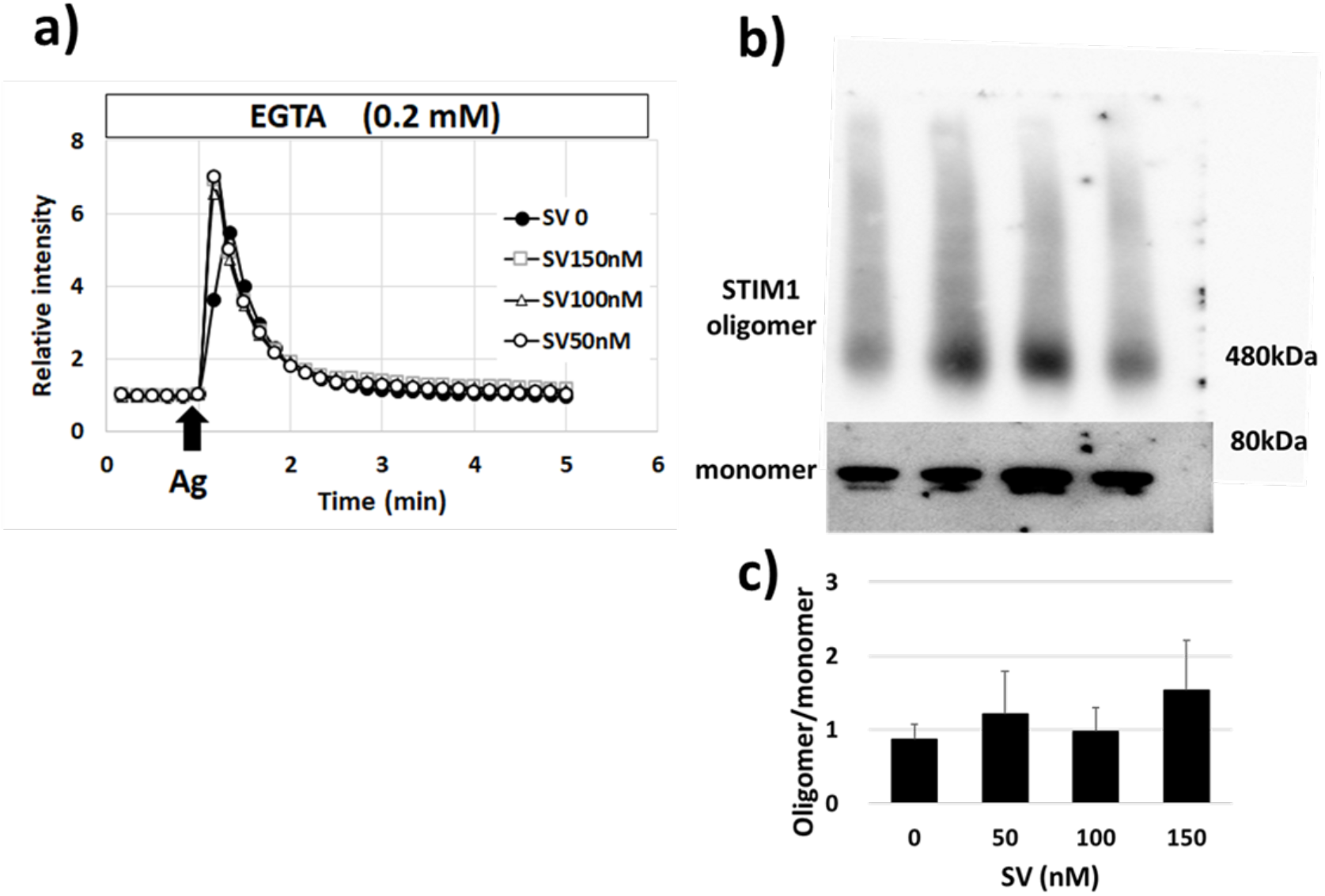
SV pretreatment does not inhibit the Ca^2+^ efflux from ER and STIM1 aggregation level by Ag-Ab stimulation. a) RBL-2H3 cells pretreated with SV for 24 h were loaded with 1 μM Fluo8/AM for 30 min and then stimulated with DNP-BSA (20 ng/mL) in calcium (-) Siraganian buffer. The change in fluorescence over time was detected using a fluorescence microscope (BIOREVO 9000, Keyence, Japan). The closed circle, open circle, open triangle, and open square lines represent SV at 0, 50, 100 nM, and 150 nM, respectively. b) Expression levels of aggregated STIM1 and total STIM1 in RBL-2H3 cells stimulated with Ag-Ab were detected using anti-STIM1 antibodies. c) Immunoreactive protein bands in B were detected using an ImageQuant™ LAS 4000 mini (Cytiva, USA) and analyzed using the ImageJ software. The statistical data show the average of the aggregated STIM1/total STIM1±SE. N=3 independent experiments.

STIM1 (stromal interaction molecule 1) is an ER membrane protein that has an EF hand motif and aggregates as an ER calcium depression following interaction with ORAI1 and the influx of extracellular Ca^2+^. We investigated whether the SV pre-treatment affected STIM1 aggregation after Ag-Ab stimulation and found that it did not have an effect (Fig. 3b,c).

These results indicate that SV does not affect Ca^2+^ efflux from the ER or influx from extracellular Ca^2+^. In addition, SV does not affect the initial signal transduction via LAT.

### Effects of SV on mast cell degranulation and counteraction by mevalonic acid and geranylgeraniol (but not farnesol)

Activation of PKCs by the influx of intracellular Ca^2+^ leads to the degranulation process, which involves small GTPases. Statins decrease the isoprenylation of small GTPases and alter their translocation. Next, we examined the effect of the mevalonate pathway intermediates mevalonate, farnesol, and geranylgeraniol.

The inhibition of degranulation by SV at 150 μM was completely counteracted in a concentration-dependent manner by co-treatment with mevalonic acid (10–200 μM) (Fig. 4a). Moreover, mevalonate and geranylgeraniol completely rescued the reduction in degranulation by Ag-Ab, Tg, or A23187 stimulation, while farnesol id not (Fig. 4b-d).

**Fig. 4.**
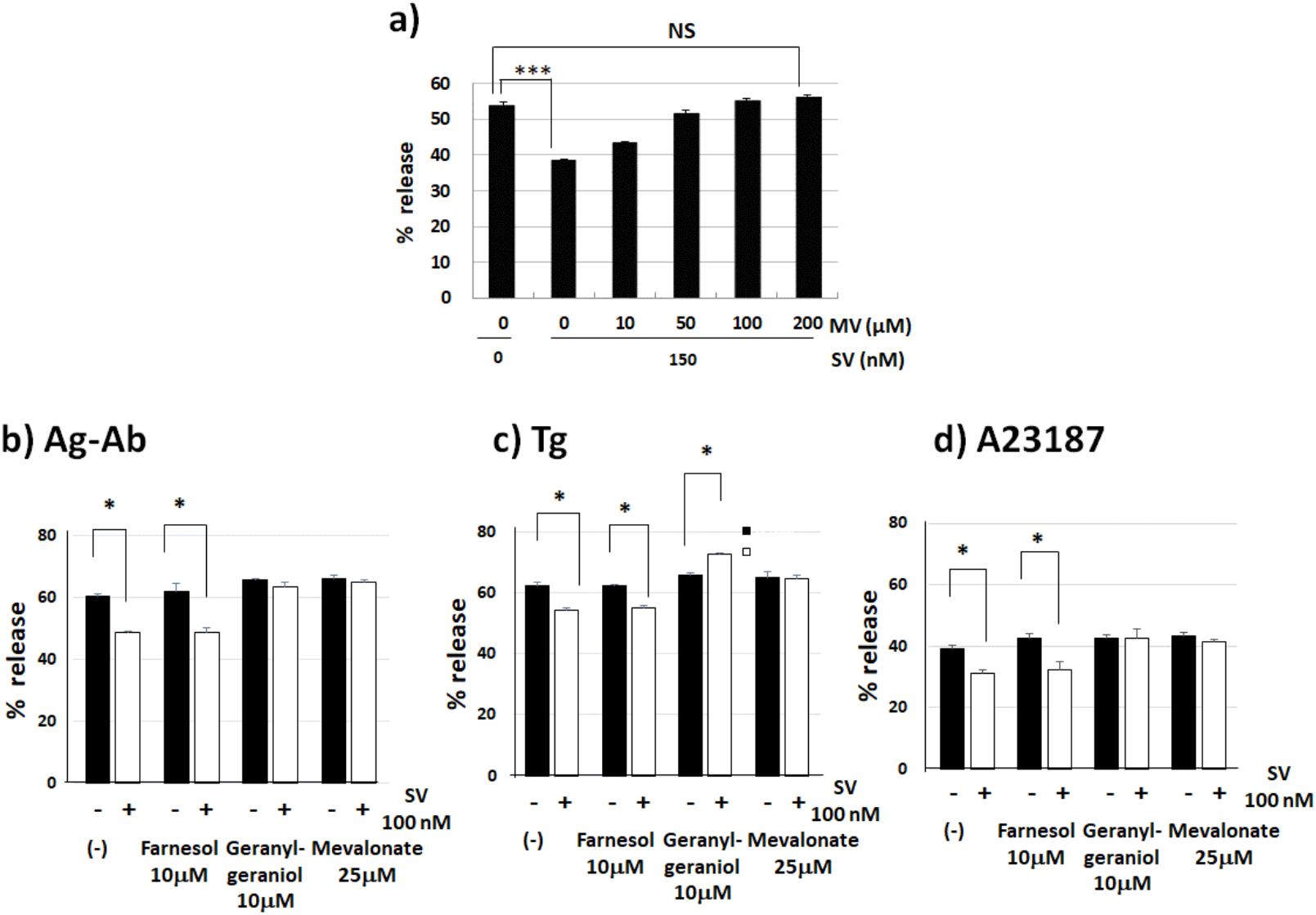
Inhibition of degranulation by SV is rescued with mevalonic acid or geranylgeraniol addition. a) IgE-sensitized RBL-2H3 cells were pretreated with 150 nM SV and 10-200 μM mevalonate for 24 h. Cells were then stimulated with DNP-BSA for 15 min. (b-d) IgE-sensitized RBL-2H3 cells were pretreated with 20 μM SV and 25 μM mevalonate, 10 μM geranylgeraniol, or 10 μM farnesol for 24 h. Cells were then stimulated with DNP-BSA (b), 1 μM Tg (c) or 500 nM A23187 (d) for 15 min. All data are presented as the mean ±SE (n=3), * p<0.05, **p<0.01(ANOVA).

### Effects of SV on PKC delta activation

SV inhibits degranulation by Ag-Ab stimulation to a greater extent compared with Tg or A23187 stimulation. Next, we tested the membrane PKC delta because PKC delta is an active regulator of mast cell degranulation induced by Ag-Ab alone. SV decreases membrane PKC delta activated by Ag-Ab stimulation but does not change the membrane PKC delta activated by Tg or A23187 stimulation (Fig. 5a-c). This difference in activation probably caused the difference in the level of degranulation between these stimulations. We examined whether this difference caused the inactivation of PKC delta by stimulating RBL-2H3 cells with Ag-Ab and the PKC activator PMA at 20 nM and found that PMA increased the degranulation rate following Ag-Ab stimulation (Fig. 5d).

**Fig. 5.**
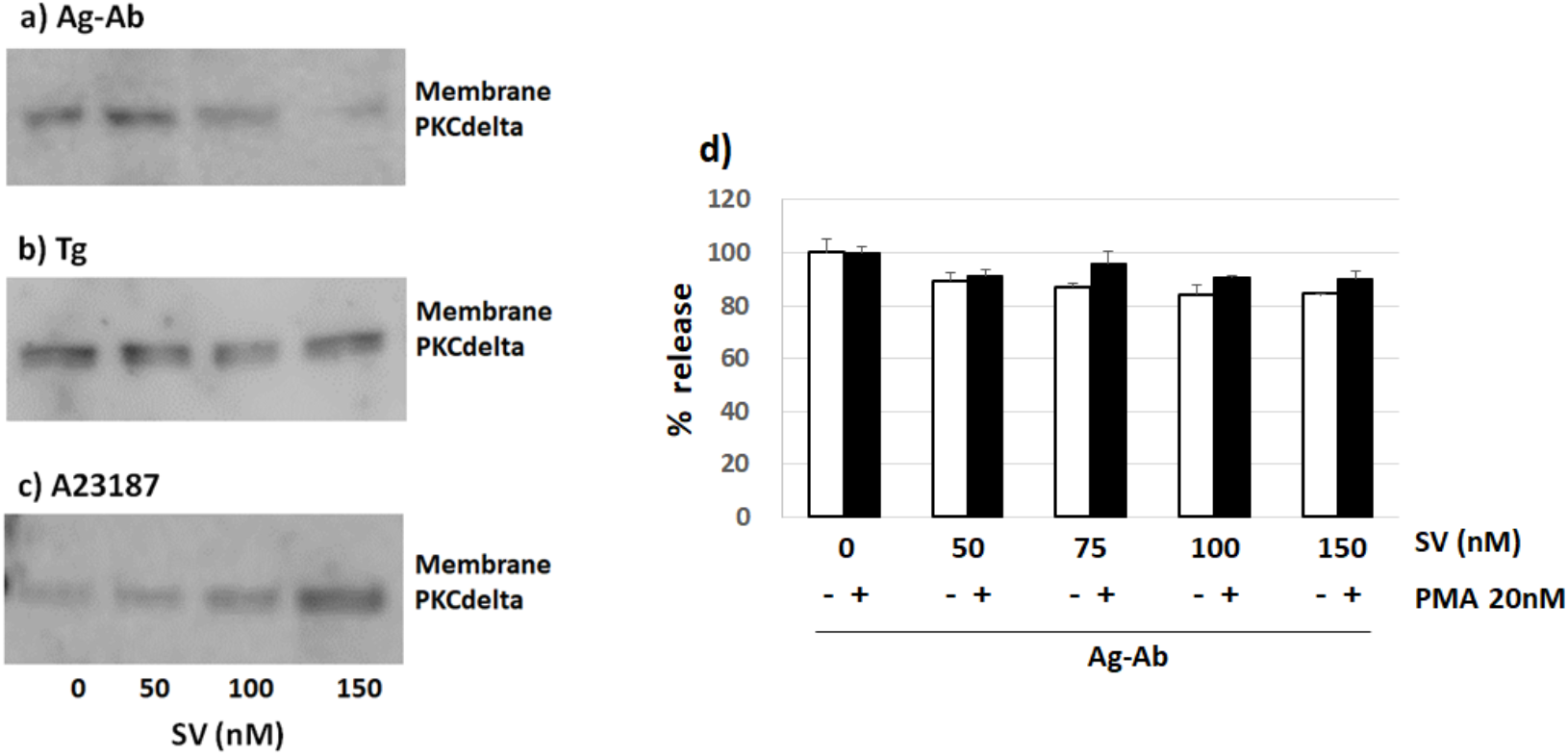
SV decreases PKC delta activation by Ag-Ab induction but not by Tg and A23187 induction. a-c) RBL-2H3 cells were incubated with SV (0, 50,100, 150 nmol/L) for 24 h at 37 °C before stimulation with Ag-Ab (Ab: cells were sensitized by DNP-IgE for 24 h, Ag: DNP-BSA 20 ng/mL, 15 min), Tg (1 μM, 15min), or A23187 (500 nM, 15 min). After stimulation, the cells were processed for SDS–PAGE and Western blot analysis as described in the text. d) RBL-2H3 cells were pretreated with DNP-IgE and SV for 24 h and stimulated for 15 min with DNP-BSA (20 ng/mL), DNP-BSA (20 ng/mL) and PMA 20 nM in Siraganian buffer.

### Effects of SV on cell morphology of RBL-2H3 cells

We then investigated the effect of SV on RBL-2H3 cell morphology to determine whether the other small GTPase Rac, which controls actin filaments, is affected by SV. We compared the morphological changes in RBL-2H3 cells after treatment with 150 nM SV and 150 nM SV+20 μM mevalonate for 24 h. Then, we observed the apparent effects on cell shape and peripheral actin filament formation in the presence of 150 nM SV for 24 h. In the RBL-2H3 cells treated with 150 nM SV, the cell shape became rounded and peripheral actin filament formation was incomplete (Fig. 6a). Actin filament formation is regulated by Rac1, CDC42, and RhoA; thus, we then investigated the degree of active Rac1. Active Rac1 was collected by pull-down assay using half of the lysates from the samples. The collected samples and remaining half of the lysates were analyzed by SDS-PAGE and detected by western blotting. In the RBL-2H3 cells treated with 150 nM SV, active Rac1 was decreased compared to that in the untreated cells (Fig. 6b).

**Fig. 6.**
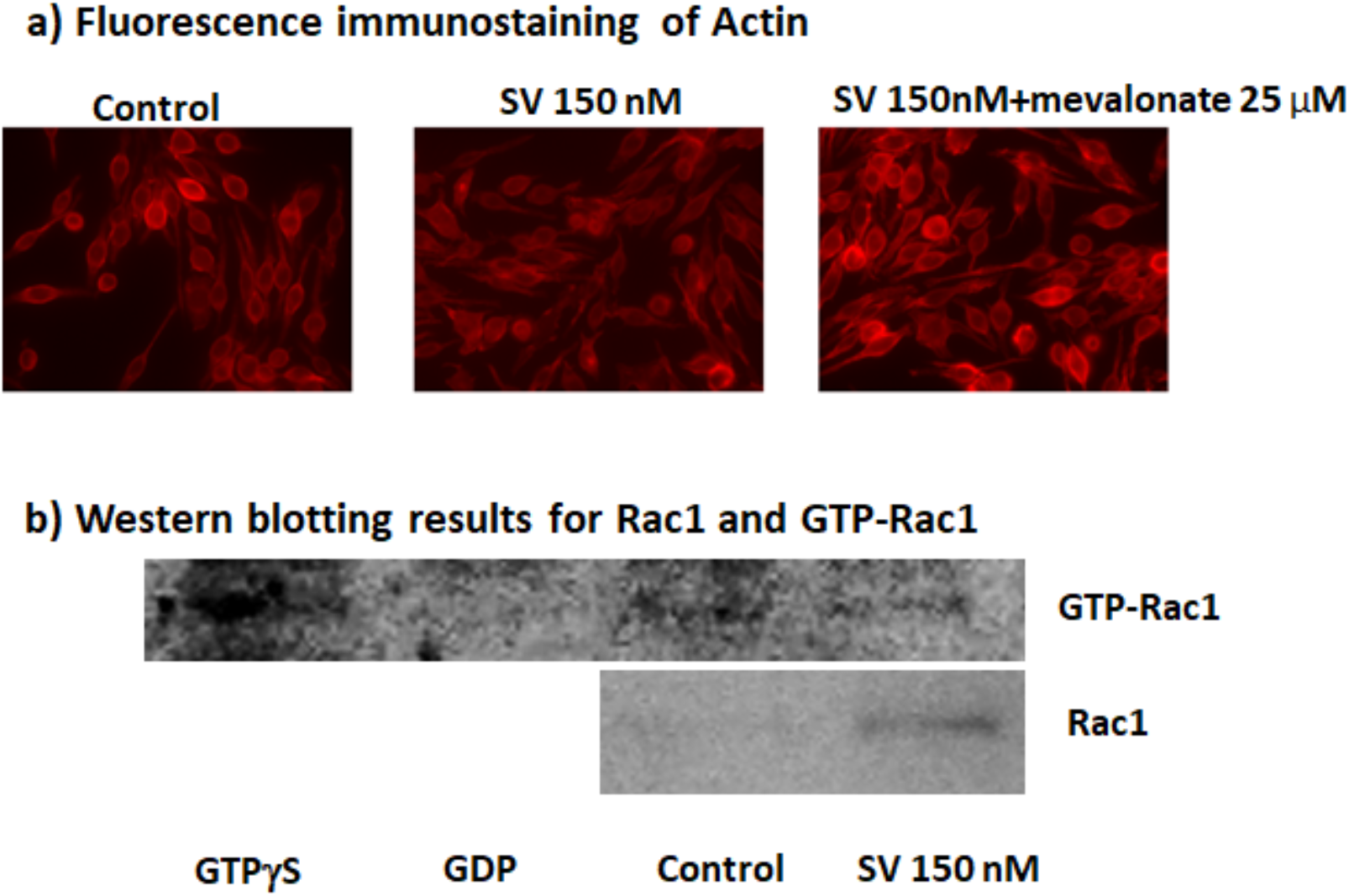
SV changes active Rac1, actin polymerization and cell morphology. a) RBL-2H3 cells were incubated with SV (150 nM) and SV (150) nM+mevalovnate (200 μM) for 24 h at 37 °C, fixed with 4% paraformaldehyde PBS, permeabilized with 70% ethanol, and incubated with phalloidin rhodamine. Images were captured using a fluorescence imaging microscope (BIOREVO 9000). b) RBL-2H3 cells were incubated with SV (150 nM) or control for 24 h at 37 °C. As positive and negative controls, cell lysates were treated with either GTPγS or GDP. The treated lysates were incubated with 400 μg GST-Rhotekin-RBD pull-down assay for active Rac1 and processed for SDS–PAGE and western blot analysis, as described in the text.

## Discussion

Statins are inhibitors of HMG-CoA reductase and represent first-choice drugs for hypercholesterolemia to reduce serum cholesterol and prevent CVD. Statins have also been reported to exert anti-inflammatory effects based on their pleiotropic characteristic. In this study, we investigated the effects of SV on the activation of RBL-2H3 cells, which is a mast cell model, because arteriosclerotic plaques include mast cells involved in the development of CVD. Fujimoto et al. ^24)^ reported the inhibitory effects of statins on histamine release from RBL-2H3 cells and found that fluvastatin had the strongest inhibitory effect on degranulation by Ag-Ab stimulation but did not affect the cytoplasmic Ca^2+^ increase or the granule content and proliferation of mast cells.

Shahid et al. ^10)^ also reported that SV (0.1~1 μM) pretreatment for 24 h inhibited histamine release from RBL-2H3 cells stimulated by Ag-Ab, ionomycin, or Tg but did not affect Ca^2+^ influx induced by these stimulations. Lytton et al. and Morgan et al. also reported that SV clearly inhibited histamine release induced by ionomycin or thapsigargin, which directly increased intracellular Ca^2+^ ^25,26)^. In another study, cerivastatin and atorvastatin did not inhibit histamine release induced by A23187 in human mast cells ^27)^.

Our data showed that SV pretreatment inhibited degranulation stimulated by Ag-Ab, Tg, or A23187 (Fig. 1) without increasing intracellular Ca^2+^ and aggregating STIM1(Fig. 2,3). The inhibitory effect of SV on degranulation by Ag-Ab stimulation was stronger than that on degranulation by Tg and A23187 stimulation.

These contradictory results could be attributed to the use of different statin types and concentrations, which were higher than the clinical range. We tested lower concentrations (≤ 150 nM) as well. The maximal clinical serum concentration was 36±19 nM after a single administration of 20 mg SV; however, SV is a hydrophobic stain (oil-water distribution coefficient of 39.8), and its concentration in plaque is higher than that in serum after clinical administration. Thus, the SV concentration in the plaque and mast cells may have reached 100 nM, which indicates that our data are consistent with the clinical condition.

Fujimoto et al. reported that the fluvastatin-induced inhibitory effect may be mediated by defects in geranylgeranyl pyrophosphate synthesis because degranulation is inhibited by the geranylgeranylation inhibitor GGTI-286 ^28)^ but not by the farnesylation inhibitor FPTIII ^29,30)^. Shahid et al. focused on three steps following the early cell signaling step: granule translocation, granule docking/priming, and granule fusion. These degranulation steps involve two processes: a Ca^2+^-dependent process for granule-membrane fusion and a Ca^2+^-independent process for microtubule formation ^31)^. Azouz et al. identified 30 Rab GTPases as potential regulators of exocytosis in RBL cells ^10)^. These Rabs split into three groups: the first group exclusively affects IgE-mediated secretion mainly mediated by kiss-and-run exocytosis ^32)^, the second group selectively affects the secretion triggered by the combination of a Ca^2+^ ionophore and the phorbol ester, which involves full exocytosis ^32)^, and the third group affects both types of secretion triggered by either stimulus ^10)^.

We also showed that co-treatment with mevalonate or geranylgeraniol counteracted the SV inhibitory effect on degranulation by stimulation; however, co-treatment with farnesol did not counteract the inhibition shown in Fig. 4. This result is consistent with that of a previous study using an inhibitor of geranylgeranyl transferase, GGTI-286 ^33)^. Sahida et al. concluded that inhibition of the mevalonate pathway by SV decreased following geranylgeranylation of Rab proteins owing to the depletion of geranylgeranyl pyrophosphate, which disrupted the interaction between Rab27a and Doc2a in the exocytosis process of RBL-2H3 cells ^12)^. Higashino et al. reported that Rab37 is a Munc13-4-binding protein that inhibits mast cell degranulation through its effector protein by counteracting the vesicle-priming activity of the Rab37-Munc13-4 system ^34)^. SV decreases geranylgeranylation of Rab proteins related to exocytotic pathways, interrupts localization of geranylgeranylated proteins, and has anti-inflammatory effects in mast cells.

As shown in Fig. 1, the SV inhibitory effect on the degranulation of RBL-2H3 cells by Ag-Ab stimulation was stronger than that of Tg and A23187 stimulation. As shown in Fig. 5, SV pretreatment inhibited the activation of PKC delta by Ag-Ab. The Ag-Ab reaction induces the activation of PLCg and PI3K, while Tg and A23187 induce PLD activation. The difference in activated signaling molecules probably caused this difference in PKC delta activation. PI3K directly activates PKC delta and PLCg hydrolyzes PIP2 to IP3 and DAG. Moreover, PLD hydrolyzes structural phospholipids, such as phosphatidylcholine, to the membrane-bound lipid phosphatidic acid (PA) and choline, while PA is converted to other lipids, such as DAG. PLD has more substrates hydrolyzed to DAG than PLC, and it is possible that PLD escapes the influence of SV on the plasma membrane. The SV inhibitory effect on the degranulation of RBL-2H3 cells by Ag-Ab stimulation with PMA was lower than that by Ag-Ab stimulation (Fig. 5b).

SV inhibited F-actin formation and decreased the activation of the Rho family Rac1 small GTPase (Fig. 6). This inhibition may be caused by a decrease in Rho family small GTPase geranylgeranylation. However, whether exocytosis of mast cells is relevant to actin rearrangement remains controversial ^20)^. Wo L-G. et al. reviewed recent progress regarding the crucial roles of F-actin in regulating exocytosis from studies on secretory cells, particularly neurons and endocrine cells. By enhancing the plasma membrane tension, F-actin promotes fusion pore expansion, vesicular content release, and a fusion mode called shrink fusion, which involves fusing vesicle shrinking ^35)^. The Rho and Ras families are involved in PI3K activity, and the Rho family is also modified by geranylgeranylation for translocation ^36)^.

In this study, we investigated the effects of SV on the degranulation of RBL-2H3 cells stimulated with Ag-Ab, Tg, or A23187. The inhibitory effect of SV is derived from the depletion of intracellular mevalonic acid and geranylgeraniol pyrophosphate, as indicated by the full recovery of these inhibitory effects of SV by mevalonic acid and geranylgeraniol but not by farnesol. Moreover, SV decreased Rac1 activation, which requires Rac1 proteins to be translocated via geranylgeranylation. SV may induce vesicular transport malfunctions by inhibiting the translocation of Rab proteins. Regarding degranulation stimulated by Ag-Ab, SV decreases PKC delta activation that may be caused by the inhibited translocation of Rho and Ras proteins.

